# ReadItAndKeep: rapid decontamination of SARS-CoV-2 sequencing reads

**DOI:** 10.1101/2022.01.21.477194

**Authors:** Martin Hunt, Jeremy Swann, Bede Constantinides, Philip W Fowler, Zamin Iqbal

## Abstract

**Summary:** Viral sequence data from clinical samples frequently contain human contamination, which must be removed prior to sharing for legal and ethical reasons. To enable host read removal for SARS-CoV-2 sequencing data on low-specification laptops, we developed ReadItAndKeep, a fast lightweight tool for Illumina and nanopore data that only keeps reads matching the SARS-CoV-2 genome. Peak RAM usage is typically below 10MB, and runtime less than one minute. We show that by excluding the polyA tail from the viral reference, ReadItAndKeep prevents bleed-through of human reads, whereas mapping to the human genome lets some reads escape. We believe our test approach (including all possible reads from the human genome, human samples from each of the 26 populations in the 1000 genomes data, and a diverse set of SARS-CoV-2 genomes) will also be useful for others.

**Availability and implementation:** ReadItAndKeep is implemented in C++, released under the MIT license, and available from https://github.com/GenomePathogenAnalysisService/read-it-and-keep.

## Introduction

Since experimental isolation of viral DNA from the host is imperfect, viral sequence data is frequently contaminated with host DNA. Removal of host sequence is a first step for many analyses, and where the host is human this is essential to safeguard patient anonymity. Typical approaches [1] either map reads directly to the host genome (eg using BWA MEM [2], Bowtie2 [3]), or use a metagenomics classifier (eg Kraken2 [4]) to assign each read to a species. However in some circumstances (such as a global pandemic following a recent zoonosis) the viral genome is known and of limited diversity, opening up the possibility of positively identifying viral reads by mapping to a reference. In this paper we develop a simple tool that scans sequence data and retains only that which maps to the viral genome. By rigorously testing both theoretically and with human data from diverse global populations predating the pandemic, we are able to give convincing evidence that mapping to a modified SARS-CoV-2 reference is sufficient to guarantee removal of human data. The tool, named ReadItAndKeep, is extremely fast and requires very little RAM – typically a few MB as compared with around 10GB for methods based on mapping to the human genome. This allows read decontamination locally on a standard laptop before uploading to a shared or public server for analysis, or depositing in read archives.

## Methods

ReadItAndKeep is implemented in C++, using the API of minimap2 to match reads to a target genome. Hits from minimap2 are used without performing full alignment (equivalent to minimap2 default command line options, reporting approximate mappings).

A read is retained if it has a match that is at least 50bp or is at least 50% of the length of the read (these are default values of user-specifiable parameters). In the case of paired reads, a pair is kept if either of the reads have a suitable match. ReadItAndKeep uses the minimap2 presets “short read” or “ont” for Illumina and Oxford Nanopore Technology (ONT) reads respectively (same as command line options `-x sr` or `-x map-ont`). Retained reads are written to gzipped FASTQ file(s).

We compared ReadItAndKeep with a standard approach of removing reads matching the human genome. We benchmarked against the tool Dehumanizer (https://github.com/SamStudio8/dehumanizer), which wraps mappy/minimap2, with its recommended reference comprising the human genome GRCh38 plus decoy and HLA sequences (details in Supplementary text). We refer to this collection of references as the “human reference” throughout.

## Results

Complete benchmarking results are shown in Supplementary Table 1, summarised in Table 1, and described below.

**Table 1:** Summary of testing ReadItAndKeep (RIAK) and Dehumanizer on Human and SARS-CoV-2 reads. Percent reads retained is calculated from summing across reads from all samples in the data set. Mean run time is the mean wall clock time used across all samples in the data set.

| Data set | Samples | Total reads | Percent reads retained |  | Mean run time |  | Peak RAM (MB) |  |
| --- | --- | --- | --- | --- | --- | --- | --- | --- |
|  |  |  | Dehumanizer | RIAK | Dehumanizer | RIAK | Dehumanizer | RIAK |
| Human 75-mers | 1 | 3,182,668,318 | 0.763 | 0.0 | 5,082m | 83m | 30,378 | 6 |
| Human 150-mers | 1 | 3,181,197,226 | 0.034 | 0.0 | 7,332m | 149m | 30,376 | 6 |
| Human Illumina | 27 | 20,772,464,024 | 0.903 | 0.0 | 2,860m | 59m | 11,831 | 8 |
| Human ONT | 1 | 15,666,887 | 10.283 | 0.0 | 9,591m | 103m | 10,839 | 243 |
| SARS-CoV-2 75-mers | 1 | 29,796 | 100.0 | 100.0 | 48.0s | 0.3s | 11,305 | 7 |
| SARS-CoV-2 150-mers | 1 | 29,721 | 100.0 | 100.0 | 48.0s | 0.3s | 11,305 | 7 |
| SARS-CoV-2 Illumina | 246 | 610,451,014 | 99.994 | 99.894 | 102.4s | 49.6s | 11,330 | 9 |
| SARS-CoV-2 ONT | 189 | 30,422,462 | 100.0 | 99.992 | 52.1s | 14.3s | 8,387 | 9 |

### Evaluation of human read removal

we first checked that ReadItAndKeep should in principle remove human reads, using all 75-mers and 150-mers of the human reference as “reads”, with the target genome SARS-CoV-2 MN908947.3. Only 90,469 (0.001%) 75-mers were retained, all of which matched the 33bp poly-A tail of MN908947.3. Since this tail provides no useful information and is excluded by SARS-CoV-2 amplicon sequencing, we removed if from the viral genome for all further analysis. Using this trimmed sequence as the target, all tested k-mers were removed by ReadItAndKeep. Dehumanizer retained 0.76% of the 75bp reads, and 0.03% of the 150bp reads (Table 1).

We then measured the success at human read removal on 27 Illumina runs from the expanded 1000 genomes project [5]: the well-studied sample NA12878 plus one sample from each of the 26 populations, originating from Africa, Asia, Europe and the Americas (Supplementary Table 2). A high depth run of ONT reads from NA12878 was also tested [6]. Note that all of these samples were sequenced years before the SARS-CoV-2 virus jumped into humans, and so we assume that all reads in these datasets should be excluded. Across all these samples, ReadItAndKeep retained zero reads, but Dehumanizer kept 1.8% Illumina and 10% ONT reads (Table 1). Further investigation of the 10% showed they were heavily enriched for very low quality and repetitive reads, with multi-kb softclipped regions.

### Quantification of SARS-CoV-2 read retention

we confirmed that all 75-mers and 150-mers from the SARS-CoV-2 reference genome were retained by Dehumanizer and ReadItAndKeep. Next, a set of genetically diverse samples was collated, comprising 246 Illumina and 189 ONT sequencing runs, chosen (see Supplementary text) to maximise unique protein mutations and ensure a range of lineages as assigned by Pangolin [7]. Dehumanizer retained >99.99% of reads, and ReadItAndKeep kept >99.99% of ONT reads and 99.89% of Illumina reads (Table 1). For diagnostic purposes, those reads excluded by ReadItAndKeep were then mapped to the SARS-CoV-2 genome using Bowtie 2 [3] with the `--very-sensitive-local` option. The excluded reads were highly enriched for low quality - with either a very short match or high error rate (see Supplementary Figure 1, Supplementary Table 3). The greatest loss in mean per-base depth was 0.21% for ONT and 1.87% for Illumina (238/246 Illumina samples had mean loss <1%) (Supplementary Table 3). We conclude this loss of a tiny volume of low quality reads would not affect downstream analyses.

## Discussion

There are broadly three options for decontaminating SARS-CoV-2 datasets: exclude reads mapping to human (as done by Dehumaniser), keep reads mapping to the virus (as done by ReadItAndKeep), or do both (first map to the virus, and then exclude any of that also map to human, as is done by the COG consortium). We have shown that, by trimming the poly-A tail from the SARS-CoV-2 genome used by ReadItAndKeep, we completely remove spurious matches of human reads. Thus ReadItAndKeep offers an approach that is more reliable than just mapping to the human genome, and lighter weight (low RAM, fast) than either of the other two approaches.

We also investigated using ReadItOnKeep for Influenza A and HIV-1 samples, which are known to be significantly more diverse than SARS-CoV-2. Although all human reads were removed, the method was not effective in retaining viral reads, in extreme cases rejecting more than half. Therefore we only recommend ReadItAndKeep for viruses with low levels of diversity - our focus was SARS-CoV-2.

Finally, one challenge for implementing pathogen sequencing in healthcare systems is justifying what proportion of human reads must be removed to guarantee non-identifiability. By explicitly testing with all possible 75bp and 150bp reads in the (extended) human reference genome, and 27 human genome samples from different global ancestries, we were able to show ReadItAndKeep excluded every single human read. We hope the benchmarking approach itself will be of use, and that the speed and low resource requirements will make ReadItAndKeep of wide utility.

## Contributions

MH, ZI designed the study and wrote the paper. MH developed the software and performed analyses. BC, JS added bioconda, docker. PF collected the viral dataset.

## References

[1] Stephen J. Bush, Thomas R. Connor, Tim E.A. Peto, Derrick W. Crook, and A. SarahYR 2020 Walker. Evaluation of methods for detecting human reads in microbial sequencing datasets. Microbial Genomics, 6(7):e000393, 2020.

[2] Heng Li. Aligning sequence reads, clone sequences and assembly contigs with BWA-MEM. arXiv:1303.3997 [q-bio], May 2013. arXiv: 1303.3997.

[3] Ben Langmead and Steven L Salzberg. Fast gapped-read alignment with Bowtie 2. Nature methods, 9(4):357–359, March 2012.

[4] Derrick E. Wood, Jennifer Lu, and Ben Langmead. Improved metagenomic analysis with Kraken 2. Genome Biology, 20(1):257, November 2019.

[5] Marta Byrska-Bishop, Uday S. Evani, Xuefang Zhao, Anna O. Basile, Haley J. Abel, Allison A. Regier, Andre Corvelo, Wayne E. Clarke, Rajeeva Musunuri, Kshithija Nagulapalli, Susan Fairley, Alexi Runnels, Lara Winterkorn, Ernesto Lowy-Gallego, The Human Genome Structural Variation Consortium, Paul Flicek, Soren Germer, Harrison Brand, Ira M. Hall, Michael E. Talkowski, Giuseppe Narzisi, and Michael C. Zody. High coverage whole genome sequencing of the expanded 1000 genomes project cohort including 602 trios. bioRxiv, 2021.

[6] Miten Jain, Sergey Koren, Karen H. Miga, Josh Quick, Arthur C. Rand, Thomas A. Sasani, John R. Tyson, Andrew D. Beggs, Alexander T. Dilthey, Ian T. Fiddes, Sunir Malla, Hannah Marriott, Tom Nieto, Justin O’Grady, Hugh E. Olsen, Brent S. Pedersen, Arang Rhie, Hollian Richardson, Aaron R. Quinlan, Terrance P. Snutch, Louise Tee, Benedict Paten, Adam M. Phillippy, Jared T. Simpson, Nicholas J. Loman, and Matthew Loose. Nanopore sequencing and assembly of a human genome with ultra-long reads. Nature Biotechnology, 36(4):338–345, April 2018.

[7] Aine O’Toole, Emily Scher, Anthony Underwood, Ben Jackson, Verity Hill, John T McCrone, Rachel Colquhoun, Chris Ruis, Khalil Abu-Dahab, Ben Taylor, Corin Yeats, Louis Du Plessis, Daniel Maloney, Nathan Medd, Stephen W Attwood, David M Aanensen, Edward C Holmes, Oliver G Pybus, and Andrew Rambaut. Assignment of Epidemiological Lineages in an Emerging Pandemic Using the Pangolin Tool. Virus Evolution, page veab064, July 2021.

