## Supplementary Material for "ReadItAndKeep: rapid decontamination of SARS-CoV-2 sequencing reads"

### – supplementary information

Martin Hunt [1,2], Jeremy Swann [2], Bede Constantinides [2], Philip W Fowler [2], Zamin Iqbal [1]

[1]European Bioinformatics Institute, Cambridge, UK and
[2]Nuffield Department of Medicine, University of Oxford, Oxford, UK.

#### Contents

**1 Funding** **2**

**2 Data and accessions** **2**

**3 Software versions and command lines** **3**

**4 Analysis of rejected SARS-CoV-2 reads** **4**

### 1 Funding

Martin Hunt is funded by the National Institute for Health Research (NIHR) Health Protection Research Unit in Healthcare Associated Infections and Antimicrobial Resistance (NIHR200915), a partnership between the UK Health Security Agency (UKHSA) and the University of Oxford. The views expressed are those of the author(s) and not necessarily those of the NIHR, UKHSA or the Department of Health and Social Care. For the purpose of open access, the author has applied a Creative Commons Attribution (CC BY) licence to any Author Accepted Manuscript version arising.

### 2 Data and accessions

##### Human reference

We used the same reference sequences as those recommended on the README of the Dehumanizer github repository (`https://github.com/SamStudio8/dehumanizer`), except for using the latest HLA sequences that have become available since README was made.

The GRCh38 reference was downloaded from `https://ftp.ncbi.nlm.nih.gov/genomes/all/GCA/000/001/405/GCA_000001405.28_GRCh38.p13/GCA_000001405.28_GRCh38.p13_genomic.fna.gz`, and the decoy sequences from `ftp://ftp.ncbi.nlm.nih.gov/genomes/all/GCA/000/786/075/GCA_000786075.2_hs38d1/GCA_000786075.2_hs38d1_genomic.fna.gz`. The HLA sequences were downloaded from `https://github.com/ANHIG/IMGTHLA/archive/refs/tags/v3.46.0-alpha.tar.gz`.

The three FASTA files were combined in to a single file called `dehuman.20211018.fasta`, and then two indexes produced using the commands:

```
minimap2 -x sr -d dehuman.20211018.sr.mmi dehuman.20211018.fasta
minimap2 -x map-ont -d dehuman.20211018.map-ont.mmi dehuman.20211018.fasta
```

These two indexes were used when running Dehumanizer, the choice determined by the sequencing technology.

##### Human Reads

Version "rel7" of the ONT reads for human sample NA12878 were downloaded from `http://s3.amazonaws.com/nanopore-human-wgs/rel7/rel_7.fastq.gz`, as described in the github repository `https://github.com/nanopore-wgs-consortium/NA12878/blob/master/Genome.md`.

The file of metadata containing the human Illumina sequencing runs was `http://ftp.1000genomes.ebi.ac.uk/vol1/ftp/data_collections/1000G_2504_high_coverage/1000G_2504_high_coverage.sequence.index` (linked from the page `https://www.internationalgenome.org/data-portal/data-collection/30x-grch38`). We chose NA12878, plus the first run in the metadata file for each population (as defined in the "POPULATION" column). The run IDs and populations are provided in Supplementary Table 2.

##### SARS-CoV-2 reads

Metadata for 509,977 SARS-CoV-2 samples were downloaded from the COGUK consortium website on 2021-06-24. We selected the 455,352 samples which had been deposited in the European Nucleotide Archive (ENA) under project PRJEB37886. The FASTQ files of around half of these (230,879) were then downloaded from the ENA. The most common 50 lineages, as classified by pangoLEARN v1.2.13, were identified and to this set were added 7 lineages that were rarer in the dataset but we wished to sample, including B.1.617.2, P.1 and P.2.

Two genetically diverse datasets (one containing Illumina reads and one containing ONT reads) were created by the following iterative process for each technology. 100 samples from the first lineage were randomly chosen with replacement, as a set of manageable size from which to select samples for our final dataset. We then iteratively added five samples, from the 100, to the final dataset by each time adding the sample that results in the greatest number of mutations in the final dataset. Next, five samples from the second lineage were added using the same iterative process, chosen from 100 random samples from the second lineage. This was repeated for the remaining lineages, adding five samples (or as many samples existed with the chosen lineage) from each lineage until all 57 lineages had been considered. Since not all lineages had five or more samples, the resulting Illumina and ONT datasets contained 246 and 189 samples, respectively. The accessions are provided in Supplementary Table 3.

#### 3 Software versions and command lines

ReadItAndKeep git commit a898e0cb758b547650f3b45264f7df063a47007a was used. The command line used for Illumina was:

```
readItAndKeep --tech illumina \
  --ref_fasta target.fasta \
  --reads1 reads_1.fastq.gz \
  --reads2 reads_2.fastq.gz \
  -o out
```

and for ONT:

```
readItAndKeep --tech ont \
  --ref_fasta target.fasta \
  --reads reads.fastq.gz \
  -o out
```

where `target.fasta` is the target genome, ie the SARS-CoV-2 reference genome.

Dehumanizer version 0.9.0 (git commit fc29e0a958c936f0f2bce59317a11eb3cd19dd5d) was used, with the reference genome file `dehuman.20211018.fasta`, described above. The command line used was:

```
dehumanise manifest.tsv \
  --preset PRESET \
  --fastx reads.fastq.gz \
  -o out.fastq \
  --log log.txt
```

where `PRESET` was either `sr` or `map-ont` as appropriate.

All run times and RAM usage were determined by prefixing the above commands with `/usr/bin/time -v`, and using the resulting “Elapsed (wall clock) time” for the wall clock time, and “Maximum resident set size (kbytes)” for the maximum RAM. For Dehumanizer on Illumina data, where one run was needed for each of the two FASTQ files, the sum of the wall clock times and maximum RAM from the two runs was used.

### 4 Analysis of rejected SARS-CoV-2 reads

All kept, and all rejected reads from ReadItAndKeep were mapped to the SARS-CoV-2 genome separately using Bowtie 2 version 2.4.4 with the option `--very-sensitive-local`. The "error rate" reported by `samtools stats` was taken to be the error rate of the reads, which divides the number of mismatches by the total number of mapped bases. The alignment length of each read was obtained from the `query_alignment_length` as reported by pysam (https://github.com/pysam-developers/pysam). Further, the read depth of the kept and rejected reads was calculated using `samtools depth -aa`. These results are provided in Supplementary Table 3.

Supplementary Figure 1 shows scatter plots of the error rates of the kept compared to rejected reads, and the proportion of aligned reads where the alignment is $< 50$bp compared to the error rate of the rejected reads. They show that for Illumina, rejected reads in general had a high error rate or a short alignment. ONT reads were rejected due to high error rates, not short alignment lengths. Therefore the rejected reads can be considered low quality and not useful for assembly and/or variant calling.

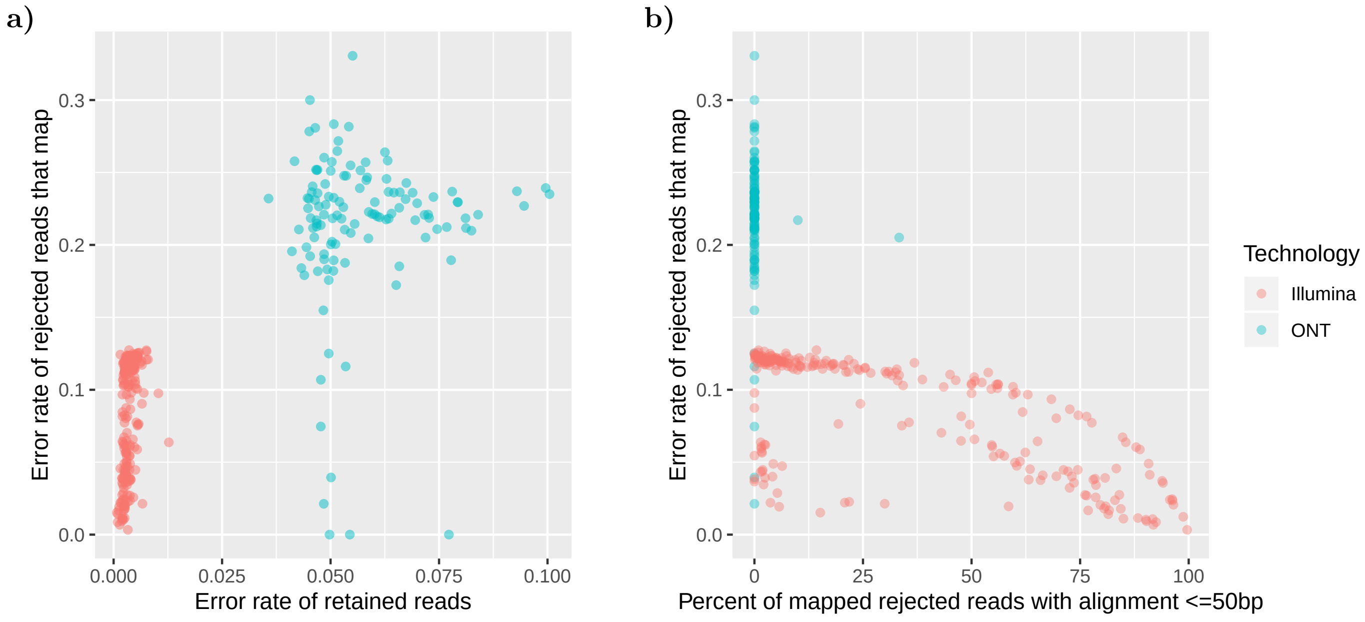


**Supplementary Figure 1:** Comparison of error rates of SARS-CoV-2 reads rejected by ReadItAndKeep, compared to: a) error rate of retained reads; b) percent of mapped reads that only have an alignment length $< 50$bp.
